# Assessment of *Culicidae* collection methods for xenomonitoring lymphatic filariasis in malaria co-infection context in Burkina Faso

**DOI:** 10.1101/2022.04.26.489492

**Authors:** Sanata Coulibaly, Simon P. Sawadogo, Achille S. Nikièma, Aristide S. Hien, Rabila Bamogo, Lassane Koala, Ibrahim Sangaré, Roland W. Bougma, Benjamin Koudou, Florence Fournet, Georges A. Ouedraogo, Roch K. Dabiré

## Abstract

**Background:** Entomological surveillance of lymphatic filariasis (LF) and malaria infections play an important role in the decision-making of national programs, to control or eliminate these both diseases. In order to corroborate infections in the human population, sampling large numbers of mosquitoes is necessary. To overcome this challenge, this study was design to assess the efficiency of four mosquito collection methods for monitoring LF and malaria infections in mosquito population.

**Methodology/Principle Findings:** Mosquito collections were performed between August and September 2018 in four villages (Koulpissy, Seiga, and Péribgan, Saptan), distributed in East and South-West health regions of Burkina Faso. Different collection methods were used: Human Landing Catches (HLC) executed indoor and outdoor, Window Exit-Trap, Double Net Trap (DNT) and Pyrethrum Spray Catches (PSC). Molecular analyses were performed to identify *Anopheles gambiae s*.*l*. sibling species and to detect *Wuchereria bancrofti* and *Plasmodium falciparum* infection in mosquitoes. A total of 3,322 mosquitoes were collected among this, *Anopheles gambiaes*.*l*. was the vector caught in largest proportion (63.82%). *An. gambiae s*.*l*. sibling species molecular characterization showed that *Anopheles gambiae* was the dominant specie in all health regions. The Human Landing Catches (indoor and outdoor) collected the highest proportion of mosquitoes (between 61.5%and 82.79%). For sampling vectors infected to *W. bancrofti* and *P. falciparum*, PSC, HLC and Window Exit-Trap were been find as the most effective collection methods.

**Conclusions/Significance:** This study revealed that HLC indoor and outdoor remained the most effective collection methods. Likewise, the results showed the probability to use Window Exit-Trap and PSC collection methods to sample *Anopheles* infected and can be useful for xenomonitoring for both LF and malaria.

**Author summary:** In Burkina Faso the monitoring and evaluation scheme to assess the impact of LF and malaria interventions is only focusing on parasitological tests. While nowadays, the most simple and direct measure of vector borne diseases is xenomonitoring. Thus, in order to confirm both diseases infection rate in the human population, sampling large numbers of mosquitoes is necessary. This study was undertaken in this context to assess the efficiency of four mosquito collection methods for xenomonitoring LF and malaria. Mosquito collections were performed between August and September 2018 in four villages, distributed in East and South-West health regions of Burkina Faso. Human Landing Catches (HLC), Window Exit-Trap, Double Net Trap (DNT) and Pyrethrum Spray Catches (PSC) were evaluated. The results showed that HLC remained the most effective collection method by collecting the highest number of *Anopheles* (2,388; 71.88% of total). Across the study, mosquito infection rate for *Wuchereria bancrofti* and *Plasmodium falciparum* were 0.004 and 0.13 respectively. To collect vectors infected it was found that Window Exit-Trap and PSC were efficiencies. In conclusion, HLC has shown to be appropriate for collect large number of mosquitoes. Likewise, Window Exit-Trap and PSC can be useful for xenomonitoring for both LF and malaria.

## Background

Vector-borne diseases are a major threat to human health worldwide. According to the World Health Organization (WHO), these diseases account for about 17% of the global burden of communicable diseases and are widespread in the poorest regions of the world [1]. Malaria and lymphatic filariasis (LF) cited as one of the main mosquitoes-borne human diseases are responsible for many cases of mortality and morbidity respectively in Sub-Saharan Africa. These are parasitic diseases which *Wuchereria bancrofti* and *Plasmodium falciparum* were responsible for most cases of LF and malaria respectively[2]. The parasites of both diseases are in majority transmitted by mosquitoes of the genus *Anopheles* in West Africa[3,4].

In this part of the world, more specifically in Burkina Faso, nearly 80.5% of cases and an estimated 4,144 deaths due to malaria were recorded throughout the country in 2017 [5]. At the same time, national neglected tropical diseases control program (NNTDCP) report revealed, that LF transmission is interrupted in 60 of the country’s 70 health districts. However, microfilaria prevalence remains above 1% in some health districts distributed in the Central, Central-Eastern, Eastern and South-Western health regions[6].

While significant progress have been made in the purview of control and elimination of LF and malaria[7–9], through vector control and chemoprevention, an effective assessment of interventions is necessary to assess the interruption of both diseases transmission. In Burkina Faso the monitoring and evaluation scheme to assess the impact of LF intervention is only focusing on parasitological tests by microfilariae diagnostic in human population but does not include the detection of parasite in mosquitoes[10]. With regard malaria control, in punctual studies, the scheme to assess the impact of intervention is done in one of two ways. First, human blood is tested for the presence of the parasite [11]. Second, mosquitoes is collected and tested, either through dissection to find the parasite [12], or through the use of molecular methods to detect the DNA [13]. Nowadays, the most direct and simple timely measure of vector borne diseases transmission is through the examination of vectors for the presence of infective stages of the parasites responsible for the infection [3]. To this effect, determine the presence of parasites in vectors, remains an option to be considered for monitoring of vector densities and their behavior in parasites transmission after setting up the control methods[14–16].

Molecular xenomonitoring, molecular method, such as polymerase chain reaction (PCR), to detect pathogen DNA or RNA in the vector is use as a proxy for infection in the human population. It has previously been used for diseases surveillance [18] after cessation of intervention [17,18] and to identify residual foci of transmission. However, in order to document that transmission has been interrupted, it is necessary to screen large numbers of vectors to ensure that the infectivity rate of vectors has reached a relatively low threshold, below which the parasite population cannot be sustained[19,20]. To do this, effective collection methods must be used for sampling potential vectors of pathogens.

The distribution and the ecological and biological characteristics of mosquitoes differ widely. Thus, of a genus to another, differences in biting and feeding behavior, resting and breeding preferences, seasonal abundance and affinity to human habitations determine their transmission potential [21]. Understanding these characteristics has enabled to set up vector collection tools such as: hand catch with oral or resting, pyrethrum spray catches, human landing catches, attractant traps, gravid traps, entry–exit trap etc. In Burkina Faso context, sampling mosquitoes relies almost exclusively upon human landing catches, which is ethically questionable due to the exposure of the collectors to infections [21,22]. Thus, alternative traps [23,24] have been compared to HLC. However, no study has compared the performance of mosquito traps for malaria and LF monitoring simultaneously in the country. Therefore, there is a need to undertake entomological surveys using diverse collection methods to assess mosquito most effective trapping methods to use for the xenomonitoring of these both diseases. To do this, the present study was undertaken to assess the efficacy of four vectors collection methods in malaria and LF surveillance in two health regions of Burkina Faso.

## Methods

### Study sites

This study was conducted in the Fada health district (East health region) likewise in Gaoua and Diébougou health districts (South-West health region). We selected four villages for the entomological surveillance: Seiga village (−0.085971°W; 11.965555°N) and Koulpissy village (−0.097974°W; 12.078119°N) located in East region, Saptan village (−3.404237°W; 1083015°N) and Péribgan village (−3.3387°W; 102218°N) located in South-West region (**Figure 1**).

**Figure 1:**
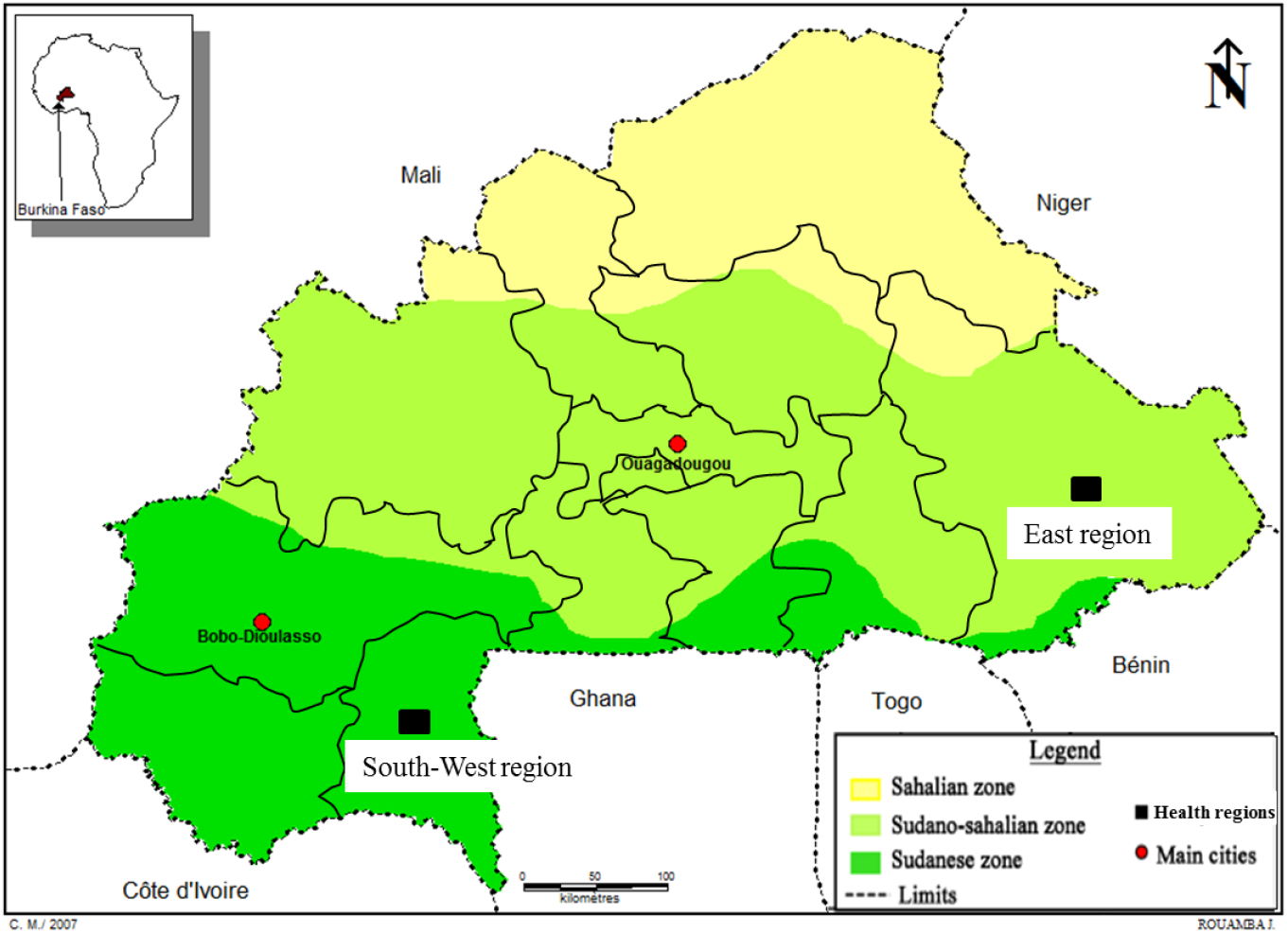
Map showing the study regions

The East region is located in the south eastern part of the country and bordered to south and east by Togo, Benin and Niger. Inside the country, the region is bordered by the region of the Centre-East, the region of Sahel and the region of Centre-North. East region is characterized by a sparse hydrographic networks and savanna type vegetation. *Anopheles gambiae s*.*l*. is the major vector of LF and malaria in this area [25].

The South-West region is located in the south of Burkina Faso. The region is bordered by Côte d’Ivoire in the south and the region of Centre-West and Ghana to the east. By the north, the region of Boucle du Mouhoun and Haut Bassins bordered the region. To the east, the South-West is bordered by the Cascades Region. South-West region is characterized by a dense hydrographic network and wooded type savannah vegetation dotted with clear forest and gallery. The major vectors of malaria and LF in the South-West region are *An. gambiae s*.*l*. and *An. funestus s*.*l*.[13,26]. A recent study has showed that *An. nili* was *W. bancrofti* potential vector [27] in this region. Other *Anopheles* species in particular *An. pharoensis, An. rufipes, An. coustani, An. flavicosta* and *An. pretoriensis* can be encountered [13].

### Collection methods

#### Human Landing Catches (HLC)

it is method which enables to sample mosquitoes seeking a human host for taking the blood feeding. Thus, the mosquitoes are collected when they land on exposed legs. This method is useful for assessing human-vector contact, host attractiveness, mosquito survival and infection and infectivity rates [28]. HLC is the most common method of collecting large number of mosquitoes, but it is ethically questionable due to the exposure of the collectors[29–31].

#### Window Exit-Trap

These traps are rectangular boxes made of a wooden or wire frame on which is stretched a mesh of braided glass fibers covered with Teflon®. On one side there is an inclined rectangular slot made of wire to allow mosquitoes to enter and on the other side there is an opening in which a cotton suction sleeve is inserted and can be closed [29,30]. Window Exit-Trap is used to monitoring some vectors species that tend to enter houses at night bite and leave the house soon after feeding without resting indoor. It provides information about exophilic versus endophilic resting behavior of vectors, physiological and biodemographic status distributions of the specimens sampled[32].

#### Double Net Trap (DNT)

It is two box nets; the inner net protects the human-bait, and the outer net is raised of the ground so that mosquitoes lured to the human-bait are collected between the nets. The nets are not treated with any insecticide [29,31]. Several study have shown that DNT capture anthropophilic *Anophelinae* females with comparable parity rates and with sporozoite indices comparable to those obtained in human landing catches but are not as efficient as human landing catches [33,34].

#### Pyrethrum Spray Catches (PSC)

This method consisted of spraying the inside of houses closed with residual aerosol insecticide very early in the morning[30].The PSC is one of the most common methods for sampling indoor-resting populations of vectors. It is used to yield information on feeding pattern survey indoor resting densities and vector species composition[32].

### Study design

This was an initial study, that is conducted in four villages (Seiga, Koulpissy, Saptan and Péribgan) distributed in East and South-West regions of Burkina Faso. Mosquitoes were collected between August and September 2018. The study was undertaken to assess the efficacy of four mosquitoes collection methods for LF and malaria monitoring in areas were malaria is endemic and LF persist. In each village, two collection days was investigated and 11 households in which mosquito samples were conducted were chosen in random way. The households were spread distributed as follows: two households for human landing catches, whose four consenting adult volunteers are recruited and trained at each site to collect mosquitoes (one indoor and other outdoor), two households for expose DNT to outdoor, two households for the Window Exit-Trap outdoor exhibition and five households for the PSC.

### Mosquito Sampling

Vectors sampling by HLC and DNT were performed in four households, from 08 pm to 06 am. These two sampling methods are carried out alternately between the concessions during the two days of collection at each site. As for the Window Exit-Trap, they were kept on windows and the vectors sampling was done from 06 am to 09 am for the two consecutive collection days. PSC were done in the morning from 06 am to 09 am.

Trapped vectors were collected with aspirator and manual collection was done for mosquitoes taken in HLC and PSC. Mosquitoes are then identified under a binocular magnifying glass using the identification key of Gillies and Coetzee [35].

Mosquito sampled were stored on silicagel in 1.5ml eppendorf tubes by species/collection method/village/period and brought to laboratory of Institut de Recherche en Science de la Santé in Bobo-Dioulasso for the molecular analyses.

### Molecular analyzes

A sample of mosquitoes collected were used for the polymerase chain reaction (PCR) analyze. The polymerase chain reaction was used to identify *An. gambiae s*.*l*. sibling species according to protocol of Santolamazza *et al*., [36] and to determine *W. bancrofti* and *P. falciparum* infection according respectively to protocol of Ramzy *et al*., [37]and Echeverry *et al*.,[38].

### Data analysis

The statistical processing of the data was done with the software R. The interface R_Studio of R version 3.3.1 (2016-06-21) was used to perform the Chi square test (*X*^*2*^) with a probability threshold *p-value = 5%* to compare the proportion of mosquitoes sampled by collection method and by health region. The infection rates of *W. bancrofti* and *Plasmodium falciparum* in mosquitoes was estimated using the Pool Screen software 2.0 (Katholi*et al*., 1995) with 95% confidence interval (CI) reported as the maximum likelihood.

### Ethics consideration

The protocol of the study received approval from the institutional ethics committee of Institut de Recherche en Science de la Santé (N°A08/2014/CEIRES). Malaria and LF prophylaxis was provided to vectors collectors.

## Results

### Mosquito abundance and composition

A total of 3,322 mosquitoes were collected in the four villages distributed in the two health regions during the study period. Morphological identification of collected mosquitoes showed that 0.54% was *An. funestus* group, 1.26% were *An. sp*, 3.19%were *Aedes sp*, 12.74% were *An. nili*, 18.45% were *Culex sp* and 63.82% were *An. gambiae s*.*l*. (**Table 1**).There was significant difference in the species composition across the four villages (*X*^*2*^ = 643.19, *df* = 7, *p-value*< 2.2e-16) distributed in two health regions. Out of the collected mosquitoes, 2,603 (78.35%) were filarial and malaria vectors belonging to members of the *An. gambiae* complex, *An. funestus* group and *An. nili*. Human Landing Catches collected largest number of mosquito 1,046 (61.5%) in East health region and 1,342(82.79%) in South-West health region (**Figure 2**).

**Table 1:** Mosquito composition by collection method in four villages distributed in two health regions of Burkina Faso (**HLC**: Human Landing Catches; **PSC**: Pyrethrum Spray Catches; **WET**: Window Exit-Trap; **DNT**: Double Net Trap).

| Mosquito subfamily | Mosquito species | Eastern health region |  |  |  |  |  |  |  | Southwestern Health Region |  |  |  |  |  |  |  | Total by mosquito specie (%) |
| --- | --- | --- | --- | --- | --- | --- | --- | --- | --- | --- | --- | --- | --- | --- | --- | --- | --- | --- |
|  |  | Seiga |  |  |  | Koulpissy |  |  |  | Péribgan |  |  |  | Saptan |  |  |  |  |
|  |  | HLC | PSC | WET | DNT | HLC | PSC | WET | DNT | HLC | PSC | WET | DNT | HLC | PSC | WET | DNT |  |
| Anophelinae | An. gambiaes.l. | 292 | 185 | 10 | 5 | 554 | 208 | 2 | 9 | 400 | 36 | 12 | 3 | 273 | 93 | 5 | 33 | 2120 (63.82) |
|  | An. funestuss.l. | 0 | 0 | 0 | 0 | 0 | 0 | 0 | 0 | 8 | 4 | 0 | 0 | 2 | 4 | 0 | 0 | 18 (0.54) |
|  | An. nili | 0 | 0 | 0 | 0 | 1 | 0 | 0 | 0 | 8 | 0 | 0 | 0 | 406 | 0 | 3 | 5 | 423 (12.74) |
|  | An. sp | 0 | 0 | 0 | 6 | 3 | 0 | 0 | 4 | 2 | 0 | 0 | 9 | 12 | 0 | 0 | 6 | 42 (1.26) |
| Culicinae | Aedessp | 0 | 3 | 0 | 57 | 0 | 0 | 0 | 40 | 0 | 0 | 1 | 2 | 0 | 0 | 0 | 3 | 106 (3.19) |
|  | Culexsp | 115 | 48 | 4 | 15 | 81 | 33 | 0 | 26 | 209 | 2 | 1 | 42 | 22 | 4 | 0 | 11 | 613 (18.45) |
| Total by collection method |  | 407 | 236 | 14 | 83 | 639 | 241 | 2 | 79 | 627 | 42 | 14 | 56 | 715 | 101 | 8 | 58 | 3322 (100) |

**Figure 2:**
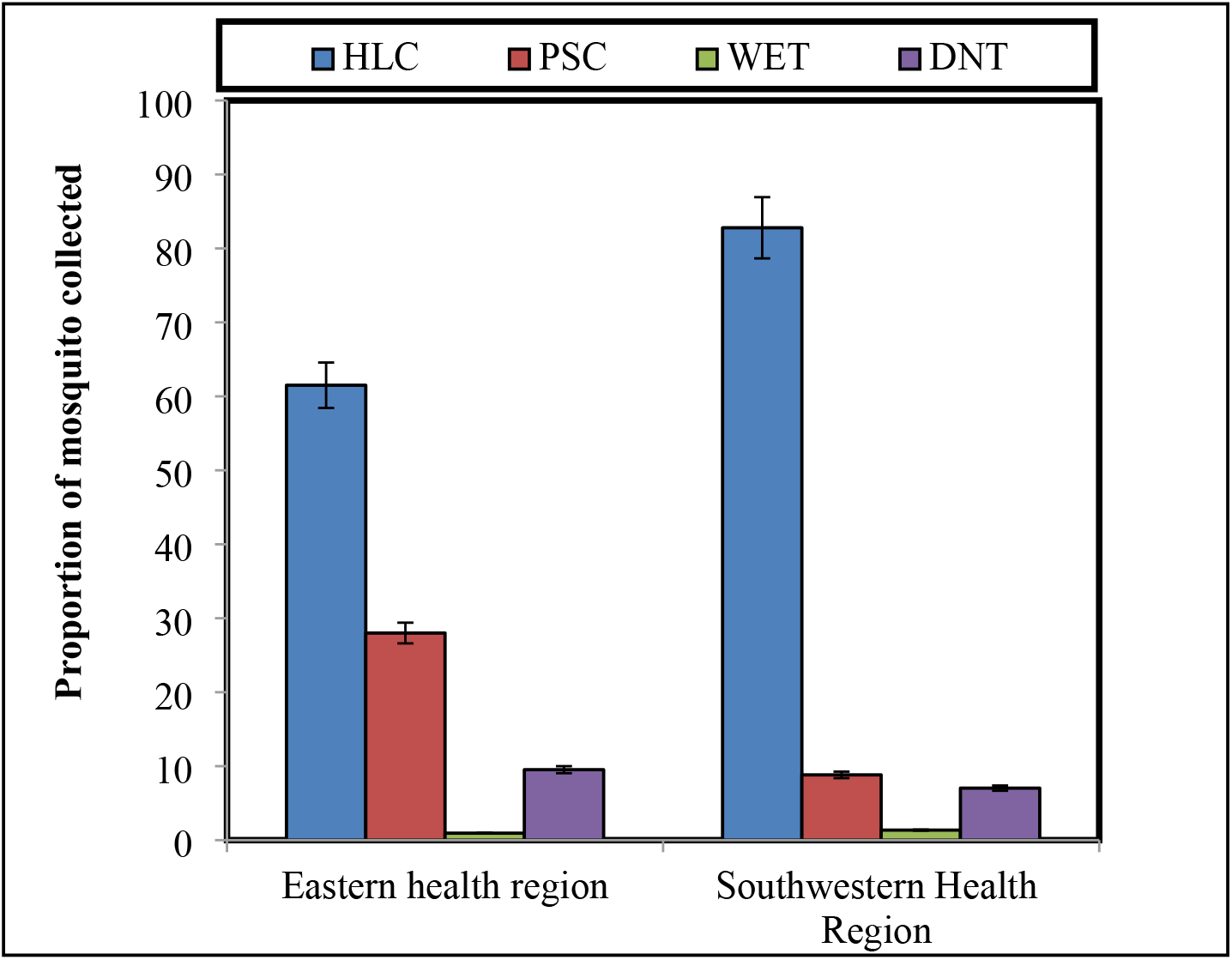
Plots of the proportion of mosquitoes caught by each method in the East and South-West health regions (*p-value< 0*.*05*) (**HLC**: Human Landing Catches; **PSC**: Pyrethrum spray catches; **WET**: Window Exit-Trap; **DNT**: Double Net Trap).

In the East region, on the 1,701 mosquitoes collected *An. gambiae s*.*l*. was collected in largest proportion using HLC (80.9%), PSC (82.4%) and Window Exit-trap. On the other hand, *Aedes sp* (47%) was collected using DNT (**Figure 3**). None *An. funestus* group has been collected in this health region.

**Figure 3:**
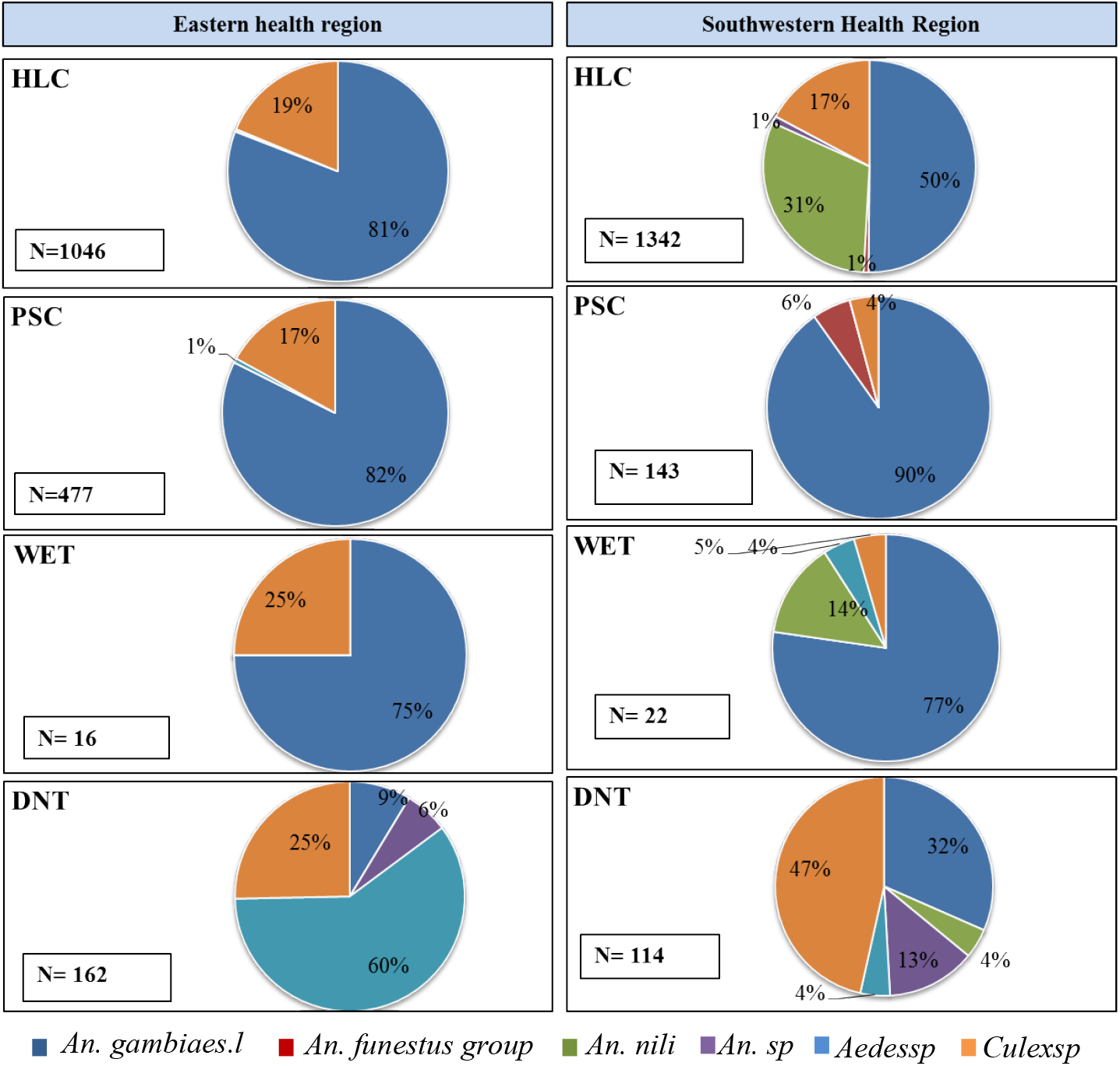
The proportion of mosquito species caught per trap type in the East and South-West health regions (Where ‘**N**’ is the number of mosquitoes caught in each trap; **HLC**: Human Landing Catches; **PSC**: Pyrethrum spray catches; **WET**: Window Exit-Trap; **DNT**: Double Net Trap).

In the South-West region, 1,621 mosquitoes were collected. *An. gambiae s*.*l*. was the mosquito specie predominantly sampled by the HLC (50%), PSC (90%) and Window Exit-trap (77%) followed by *An. nili* collected in 31% and 14% by HLC and Window Exit-Trap respectively. *Culex sp* (47%) was collected using DNT in this health region (**Figure 3**).

Molecular characterization of 327 *An. gambiae s*.*l*. sampled in the South-West region show that, 90.83% were *An. gambiae*, 7.95% were *An. coluzzii*, and 1.22% were *An. arabiensis*. In the East region, 410 *An. gambiae s*.*l*. were analyzed; 70% were characterized as *An. gambiae*, 28.54% were *An. coluzzii* and 1.46% was *An. arabiensis*. These proportions did not differ significantly between the collection methods (*X*^*2*^ = 0.8, *p-value* = 0.6).

### Mosquito infection rate

The head and thorax of mosquitoes from the *Anopheles* genera (*An*.*gambiae s*.*l*., *An. nili, An. funestus s*.*l*., *An. sp*) was analyzed for *P. falciparum* and *W. bancrofti* infection detection. Only *An. gambiae* sibling species have been found to be infected. From a total of 815 heads and thorax analyzed, sporozoid index was 0.13 and the microfilaria index was 0.004 in all regions (**Table 2**). Pyrethrum spray catches was the most effective collection method for sampling vectors infected to *P. falciparum* in the East region followed by the Window Exit-Trap in the South-West region during the vectors collection period. Regarding the mosquito microfilaria infection, one vector was identified in the South-West region sampled by the PSC collection method. In the East region two vectors were identified, one sampled by HLC indoor and the other by HLC outdoor. No mosquito analyzed was found co-infected with both *P. falciparum* and *W. bancrofti*.

**Table 2:** Sporozoid and microfilaria index in mosquitoes by collection method according to every health region (**HLC**: Human Landing Catches; **PSC**: Pyrethrum Spray Catches; **IS**: Sporozoid Index; **ImF**: microfilaria index; **TP**: total of vectors positive).

| Collection<br>method | Easternhealthregion |  |  | Southwesternhealthregion |  |  | General |  |  |
| --- | --- | --- | --- | --- | --- | --- | --- | --- | --- |
|  | Numberof<br>mosquitoesscreened | IS (TP) | ImF (TP) | Number<br>of<br>mosquitoesscreened | IS (TP) | ImF (TP) | Numberof<br>mosquitoesscreened | IS (TP) | ImF (TP) |
| HLC Indoor | 110 | 0.1 (11) | 0.009 (1) | 160 | 0.05 (8) | 0 | 270 | 0.07 (19) | 0.0037 (1) |
| HLC Outdoor | 91 | 0.08 (7) | 0.01 (1) | 137 | 0.05 (7) | 0 | 228 | 0.06 (14) | 0.0043 (1) |
| PSC | 201 | 0.28 (56) | 0 | 89 | 0.17 (15) | 0.01 (1) | 290 | 0.24 (71) | 0.0034 (1) |
| Window Exit-trap | 8 | 0.13 (1) | 0 | 10 | 0.3 (3) | 0 | 18 | 0.22 (4) | 0 |
| DNT | 7 | 0 | 0 | 2 | 0 | 0 | 9 | 0 | 0 |
| Overall | 417 | 0.18 (75) | 0.005 (2) | 398 | 0.08 (33) | 0.003 (1) | 815 | 0.13 (108) | 0.004 (3) |

## Discussion

The sustained success of vectors borne disease elimination depends, on a careful and comprehensive monitoring parasite infection in vector populations, to detect potential persistence and/or recrudescence after diseases control tools setting up, particularly in high-risk areas. However, several methods are used to collect infected mosquitoes [39], potential vectors of certain diseases such as LF, dengue, malaria, etc [40–42]. Still, there is a challenge to identify efficient vector collection methods for *Anopheles* mosquitoes, the primary vectors of LF and malaria in sub-Saharan Africa [16]. This study reports on the first evaluation of different mosquito collection methods for monitoring LF and malaria simultaneously in Burkina Faso.

In this study, *Anopheles* genus was caught in largest proportion using the HLC and PSC. Too, *An. gambiaes*.*l*. was collected mainly with a significant difference in the abundance of mosquito species sampled at each region. Indeed, the collections were carried out during the rainy season (between August and September 2018). Thus, the presence of *An. gambiae s*.*l*. in high proportion is probably linked to the presence of the species’ preferred breeding sites, which are mostly temporary sites. The potential reasons for HLC and PSC attractiveness between collection methods is probably the season in which collections were conducted. This can be considered as one of the limitations of our study, as it was carried out only once (during the rainy season).To this effect, rains presence may make some traps (DNT and Window Exit-Trap) less attractive compared to others (HLC and PSC) in the collection of *Anopheles* which are anthropophilic vectors. In the assessment of vectors collection methods, Irish *et al*.,[39] found that the rainy season can be a factor limiting the attractiveness of a trap in collecting potential vectors of LF, due to the presence of alternative favorable components for their reproduction. The molecular identification of sibling species of *An. gambiae* complex has showed that *An. gambiae* and *An. coluzzii* were the most predominant in both health regions. *An. gambiae* and *An. coluzzii* repartitions are correlated and their geographical distribution has not changed much recent years [40,41] in Burkina Faso.

In the East region, the Window Exit-Trap was the least attractive trap compared to the average number of mosquitoes collected per method. The low mosquitoes proportion catch of the trap can be explained by the resting behavior of the majority vectors because Window Exit-Trap is useful for sampling mosquitoes with exophilic behavior and to trap mosquitoes that leave houses for oviposition [29]. In our context vectors have probably endophage/exophage and endophilic behavior because, the majority was sampled by HLC and PSC.

Pyrethrum Spray Catches and Window Exit-Trap were collection methods less attractive in mosquitoes sample in the South-West region. The low mosquito catch by these traps in this region could be explained by the vectors blood feeding behavior that are probably endophage and/or exophage because HLC collection methods have sampled the highest proportion of vectors.

Double net trap was found to be the effective trap in the *Aedes sp* and *Culex sp* collection during our sampling period. The highest proportion of *Culicinae* collected with this trap has been demonstrated by previous studies [39,42] in the collection of *Culex sp* potential vectors of LF in Brazil and *Aedes sp* dengue vectors in China.

The number of samples through analyzed PCR for the searching *P. falciparum* and *W. bancrofti* gene in this study was low (as, in the surveillance phase of LF requires processing about 10 000 mosquitoes). However, they illustrate the utility of detecting parasite DNA in mosquitoes. Thus, the infected vectors obtained from the traps, supports the evidence that these methods are useful for sampling mosquitoes and to carry out pertinent monitoring of vector borne diseases in Burkina Faso. Hence, these collection methods could be employed in monitoring vector populations which can provide valuable information to support national programs decision to stop mass treatment in national level. Comparison of the differences in vectors infection index between collection methods and locations were not performed. Therefore, this is a limitation as this information is important and can inform vector monitoring campaigns. In perspective, it would be interesting to evaluate the effectiveness of different collection methods during different times of the year to sample potential vectors, to compare differences in vectors infection index between collection methods and sites, to examine the use of other collection methods such as *Anopheles* gravid trap (AGT) used in Ghana for the collection vectors potential of *P. falciparum* and *W. bancrofti*[30].

## Conclusion

Human landing catches traditional collection method was a very efficient collection method compared with the other traps in both study regions, but particularly in the East region where highest mosquitoes were found resting indoors. While the PSC, Window Exit-Trap and HLC showed efficiency in trapping infected mosquitoes, there are limitations in relation to the fact that the collection was done only once and during the rainy season. Also, the number of mosquitoes analyzed by PCR was low.

## Author’s contributions

RKD, ASH, and SPS conceived and designed the study. SC, SPS LK and ASN performed the field study, analyzed the data, and drafted the manuscript. SC and RB assured the laboratory analyses. FF, BBK, RWB, IS and AGO reviewed the first draft of the article. RKD is guarantor of the study. All authors contributed to the drafts and read and approved the final manuscript.

## Acknowledgments

The authors express their gratitude to community health workers for their availability during the study periods. They thank warmly the inhabitants of the study sites for having accepted the implementation of the study in their villages and to have gone with. They thank likewise the national program to fight against neglected tropical diseases and the national program to fight against malaria for theircollaboration.

